# LtrDetector: A modern tool-suite for detecting long terminal repeat retrotransposons de-novo on the genomic scale

**DOI:** 10.1101/448969

**Authors:** Joseph D Valencia, Hani Z Girgis

## Abstract

Long terminal repeat retrotransposons are the most abundant transposons in plants. They play important roles in alternative splicing, recombination, gene regulation, and genomic evolution. Large-scale sequencing projects for plant genomes are currently underway. Software tools are important for annotating long terminal repeat retrotransposons in these newly available genomes. However, the available tools are not very sensitive to known elements and perform inconsistently on different genomes. Some are hard to install or obsolete. They may struggle to process large plant genomes. None are concurrent or have features to support manual review of new elements. To overcome these limitations, we developed LtrDetector, which uses signal-processing techniques. LtrDetector is easy to install and use. It is not species specific. It utilizes multi-core processors available in personal computers. It is more sensitive than other tools by 14.4%–50.8% while maintaining a low false positive rate on six plant genomes.

## 1 Introduction

Formerly considered “junk DNA”, the intergenic sequences of genomes are attracting increased attention among biologists. A particularly striking feature of these regions is the prevalence of transposable elements (TEs), a type of repeated sequence. TEs include class I elements, which replicate using RNA to “copy-and-paste” themselves, and class II elements, which replicate via a “cut-and-paste” mechanism using DNA as an intermediate [1]. Barbara McLintock discovered transposons in 1950 while studying the maize genome [2]. TEs are common to all eukaryotes, comprising around 45% of the human genome and up to 80% of some plants like maize and wheat [3, 4].

TEs have several important functions.*Bennetzen and Wang* highlight the known functions of plant TEs [5]. Transposons are the major factor affecting the sizes of plant genomes [6–8]. Under stressful conditions, they can rearrange a genome [9–11]. TEs play roles in relocating genes [12, 13], and generating new genes [14,15] and new pseudo genes [16,17]. They can contribute to centromere function [18,19]. TEs can regulate the expression of nearby genes via several mechanisms including: (i) providing regulatory elements, such as promoters and enhancers, to nearby genes [14, 20–22]; (ii) inserting themselves into genes, then targeting the epigenetic regulatory system [23]; (iii) producing small interfering RNA specific to host genes [24–26]; and (iv) generating new micro RNA genes modulating host genes [27–29]. Transposons have been utilized in cloning plant genes in a technique called transposon tagging [30–32]. They also have the potential to become a new frontier in enhancing the productivity of crops [33,34].

Long Terminal Repeat retrotransposons (LTR-RTs) are a particularly interesting type of class I transposable element related to retroviruses. LTR-RTs are widespread in plants and are considered one of their primary evolutionary mechanisms [35]. *Gonzalez, et al.* summarizes some of their functions [36]. LTR-RTs can insert adjacent to and inside of genes and promote alternative splicing [37], They play roles in recombination, epigenetic control [38,39], and other forms of regulation [36]. LTR-RTs have been found with regulatory motifs that promote defense mechanisms in damaged plant tissues [40]. They can also serve as genomic markers for evolutionary phylogeny [41].

LTR-RTs are named for their characteristic direct repeat — typically 200–600 base pair (bp) long [42]. These direct repeats surround interior coding regions (the*gag* and*pol* genes). Lerat suggests 5kbp–9kbp as a size range for LTR-RTs [1].

Computational tools are extremely important in locating repeated sequences, including LTR-RTs. Tools can be roughly divided into knowledge-based tools, which leverage consensus sequence databases to search for repeats, and de-novo tools, which use sequence comparison and structural features to search for repeats without prior knowledge about the sequence [1]. Knowledge-based methods include well-known bioinformatics software such as NCBI BLAST [43] and RepeatMasker (http://www.repeatmasker.org); they can be utilized in locating all types of known TEs including LTR-RTs. However, if the sequence of the repetitive element is unknown, these two tools cannot find copies in a genome. Several methods for locating all types of TEs *de-novo* have been developed [44–47]. De-novo tools for detecting LTR-RTs include LTR STRUC [48], MGEScan-LTR [49], LTRharvest [50], LTR seq [51], and LTR Finder [52]. Postprocessing tools like LTR retriever may help increase the accuracy of de-novo approaches [53].

These tools face a variety of usability, scalability, and accuracy concerns. For example, LTR STRUC, one of the pioneering tools for locating LTR-RTs, was developed exclusively for an old version of Windows, making it difficult to use nowadays. Several tools have external dependencies which greatly complicate their installation. None of them take advantage of the concurrent architecture of modern personal computers. All may be unable to process large plant genomes such as the barley genome on an ordinary personal computer. Some tools are highly sensitive to species-specific parameters. All produce false positive predictions and do not retrieve all known LTRs. Finally, none of these tools was designed with post-processing manual review in mind.

Thousands of plant genomes are being sequenced currently and in the near future. The 10KP Project for plant genomes (https://db.cngb.org/10kp/) and the Earth Biogenome Project (https://www.earthbiogenome.org) aim at sequencing a large number of plant genomes. This expansion of genomic data creates an urgent need for modern software tools to aid in detecting LTR-RTs in the new plant genomes; such tools should remedy the limitations of the currently available tools.

To this end, we have developed LtrDetector, which is a software tool for detecting LTR-RTs. LtrDetector depends on signal processing techniques. It is easy to install because it does not have any external dependencies. It is concurrent by default, taking advantage of the advanced hardware available on personal computers. It is not species specific. It is more sensitive to known LTR-RTs than the related tools. It can process large genomes such as the barley genome. It can produce images to facilitate the manual review/annotation of the newly located LTR-RTs.

We have tested LtrDetector on synthetic sequences and multiple genomes. Our results demonstrate that LtrDetector is the best*de-novo* tool currently available for detecting LTR-RTs.

## 2 Results and discussion

### Contributions

Our efforts have resulted in the following contributions:

- The LtrDetector software for discovering LTR retrotransposons in assembled genomes. LtrDetector is available on GitHub https://github.com/TulsaBioinformaticsToolsmith/LtrDetector(http://github.com/TulsaBioinformaticsToolsmith/LtrDetector) and in Additional File 1.
- Visualization script to view scores, which should aid in the manual verification of newly found elements — available in Additional File 2 and the GitHub repository.
- Novel pipeline to generate ground truth (sequences of known LTR retrotransposons). The pipeline is available in Additional File 2 and the GitHub repository.
- Putative LTR retrotransposons of six plant genomes (Additional Files 3–9).

### Evaluation measures

True Positives (TP) are the detections that overlap with an entry in the ground truth. False Negatives (FN) are elements listed in the ground truth but not found by a tool. False Positives (FP) are the detections that overlap with an entry in the false positive data set. Mutual overlap is required to be 95%; for example, sequences A and B are counted as equivalent if the overlapping segment between A and B constitutes 95% of both A and B. Because we calculate this overlap on the whole element rather than on its flanking LTRs, it is theoretically possible that this definition of overlap may overlook some slight inaccuracies in the length of the LTRs. We use the standard measures of sensitivity (Equation 1) and precision (Equation 2) to assess the performances of LtrDetector and the related tools. Sensitivity is the ratio (or percentage) of the true elements found by a tool, whereas precision is the ratio (or percentage) of the true elements identified by a tool to the total number of regions predicted by the same tool. Additionally, we report the F1 measure (Equation 3), which combines sensitivity and precision.

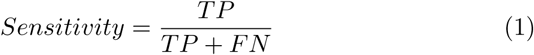

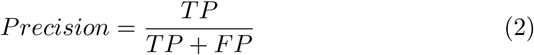

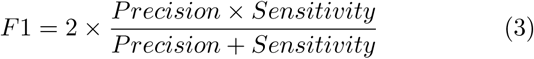

### Results on the X chromosome of the *Drosophila melanogaster*

Our initial test is based on the experiment by Lerat [1]. Table1 shows the performances of these four tools: LTR finder, LTR seq, LTRharvest, and LtrDetector. Although other tools like LTR STRUC and MGEScanLTR exist to detect LTR-RTs, they all had issues with availability and/or installation, so we were unable to get them to produce results. LtrDetector finds one fewer element that LTRharvest (91/96 vs. 92/96), while making 25% fewer total predictions (148 vs. 200). LTR seq performed the worst of the tools on every metric, and will be excluded from further experiments. These results are an early indication that LtrDetector performs well relative to the competition.*D. melanogaster* has a small genomic size and extremely well-preserved LTR sequences, making this a relatively easy test. Further evaluations are necessary to accurately gauge the performance of any tool.

**Table 1.** Results on the X Chromosome of D. melanogaster: We evaluated four de-novo tools on a ground-truth annotation provided by Lerat [1]. Total is the number of proposed LTR-RTs, TP stands for true positives.

| Tool | Total | TP (of 96) | Sensitivity | Memory(MB) | Time(s) |
| --- | --- | --- | --- | --- | --- |
| LTR_Finder | 57 | 48 | 50.0% | 300.3 | 389.7 |
| LTR_seq | 204 | 48 | 50.0% | 874.4 | 7261.7 |
| LTRharvest | 200 | 92 | 95.8% | 190.5 | 22.6 |
| LtrDetector | 148 | 91 | 94.8% | 330.5 | 92.6 |

### Results on synthetic data sets

We built synthetic data by randomly generating exact repeats — long terminal repeats — within a certain size range, mutating a selected percentage of one of them, and inserting them with random sequences in between. Although this data set does not accurately simulate the content of a real genome, it can help us demonstrate the ability of a tool to detect repeats at a given level of mutation. LTR Finder is one of the best-performing predecessor tools to LtrDetector, but its results are not listed because its strict filtering system requires other structural features that our synthetic genome lacks. For each trial, both tools are run with a sequence identity threshold of 5% lower than the similarity implied by the mutation rate. For example, the trial with 15% mutation would have an identity threshold of 80%. All other parameters are left at their defaults. Both tools capture nearly all of the elements in well conserved repeats (0–5%), but by 15% mutation, LtrDetector identifies 74 of 92 ground truth elements, whereas LTRharvest finds only 29. On the 20% mutation rate, LtrDetector outperformed LTRharvest by a wide margin in terms of the sensitivity (40/93 vs. 8/93). Neither tool is capable of reliably detecting repeats at 30% or greater mutation rates. These results are indicative of LtrDetector’s capabilities on repeats of varying levels of degeneration.

Results on six plant genomes Our main experiment was an evaluation of three tools (LtrDetector, LTR Finder and LTRharvest) on six plant genomes (including several important crops) of varying size and repeat content. All tools were run with

**Table 2.** Results on synthetic genomes: We constructed several synthetic genomes with randomly generated direct repeats mutated at a given percentage of nucleotides (0–30%) to assess performance at different levels of LTR conservation. Total is the number of proposed LTR-RTs, TP is number of true positives, GT is number of elements in the synthetic ground truth.

| Tool | Total | TP | GT |
| --- | --- | --- | --- |
| *Mutation 0%* |  |  |  |
| LTRharvest | 90 | 90 | 90 |
| LtrDetector | 90 | 90 | 90 |
| *Mutation 5%* |  |  |  |
| LTRharvest | 92 | 91 | 92 |
| LtrDetector | 90 | 90 | 92 |
| *Mutation 10%* |  |  |  |
| LTRharvest | 78 | 74 | 91 |
| LtrDetector | 88 | 88 | 91 |
| *Mutation 15%* |  |  |  |
| LTRharvest | 32 | 29 | 92 |
| LtrDetector | 74 | 74 | 92 |
| *Mutation 20%* |  |  |  |
| LTRharvest | 10 | 8 | 93 |
| LtrDetector | 40 | 40 | 93 |
| *Mutation 30%* |  |  |  |
| LTRharvest | 1 | 0 | 90 |
| LtrDetector | 2 | 2 | 90 |

**Table 3.** Results on six plant genomes: We tested three tools on one model organism, A. thaliana, and five important crops of varying genomic size and repeat content. All genomes were processed using the default parameters for each tool. We used an additional utility to process each of LTR Finder and LtrDetector concurrently because neither supports multi-threading. We did so to ensure fair comparison in terms of time since our tool, LtrDetector, is concurrent by default. Total is the number of proposed LTR-RTs, TP is number of true positives, GT is number of elements in the ground truth, FP are false positives. Sensitivity, Precision, and F1 are defined by Equations 1, 2, and 3. We report all measures for each genome and in total. Note: Results are unavailable for two tools on H. vulgare because they crashed repeatedly on this large genome. their default parameters. Results for LTRharvest and LTR Finder are unavailable for the *Hordeum vulgare* (barley) genome because both tools repeatedly caused the computer to crash. The results of this experiment suggest substantial performance gains for our tool over previous methods.

| Tool | Total | TP | GT | FP | Sensitivity | Precision | F1 | Time (hh:mm:ss) | Memory (GB) |
| --- | --- | --- | --- | --- | --- | --- | --- | --- | --- |
| *Arabidopsis thaliana* |  |  |  |  |  |  |  |  |  |
| LTR_Finder | 386 | 106 | 256 | 0 | 0.414 | 1.000 | 0.586 | 00:29:52 | 1.08 |
| LTRharvest | 1787 | 175 | 256 | 3 | 0.684 | 0.983 | 0.806 | 00:00:34 | 0.36 |
| LtrDetector | 1355 | 195 | 256 | 8 | 0.762 | 0.961 | 0.850 | 00:07:40 | 0.81 |
| *Oryza sativa* |  |  |  |  |  |  |  |  |  |
| LTR_Finder | 5061 | 1101 | 1727 | 11 | 0.638 | 0.990 | 0.776 | 04:55:20 | 1.18 |
| LTRharvest | 6684 | 1025 | 1727 | 115 | 0.594 | 0.899 | 0.715 | 00:01:42 | 0.46 |
| LtrDetector | 6559 | 1305 | 1727 | 84 | 0.756 | 0.940 | 0.838 | 00:23:51 | 1.44 |
| *Sorghum bicolor* |  |  |  |  |  |  |  |  |  |
| LTR_Finder | 11581 | 2869 | 4697 | 66 | 0.611 | 0.978 | 0.752 | 10:15:33 | 3.01 |
| LTRharvest | 17894 | 1812 | 4697 | 398 | 0.386 | 0.820 | 0.525 | 00:03:54 | 0.25 |
| LtrDetector | 21558 | 3539 | 4697 | 165 | 0.753 | 0.955 | 0.843 | 00:59:05 | 0.95 |
| *Glycine max* |  |  |  |  |  |  |  |  |  |
| LTR_Finder | 10491 | 1432 | 2876 | 6 | 0.498 | 0.996 | 0.664 | 20:19:54 | 2.37 |
| LTRharvest | 19862 | 1810 | 2876 | 4 | 0.629 | 0.998 | 0.772 | 00:06:16 | 0.63 |
| LtrDetector | 18979 | 2162 | 2876 | 12 | 0.752 | 0.994 | 0.856 | 01:03:57 | 6.86 |
| *Zea mays* |  |  |  |  |  |  |  |  |  |
| LTR_Finder | 60731 | 13414 | 20107 | 13 | 0.667 | 0.999 | 0.800 | 104:40:37 | 15.05 |
| LTRharvest | 85298 | 9536 | 20107 | 19 | 0.474 | 0.998 | 0.643 | 00:11:05 | 2.47 |
| LtrDetector | 97338 | 14326 | 20107 | 55 | 0.712 | 0.996 | 0.831 | 03:51:13 | 1.50 |
| *Hordeum vulgare* |  |  |  |  |  |  |  |  |  |
| LTR_Finder | — | — | — | — | — | — | — | — | — |
| LTRharvest | — | — | — | — | — | — | — | — | — |
| LtrDetector | 176941 | 4355 | 5792 | 172 | 0.752 | 0.962 | 0.844 | 08:57:39 | 15.93 |
| Total (Excluding *H. vulgare*) |  |  |  |  |  |  |  |  |  |
| LTR_Finder | 88250 | 18922 | 29663 | 96 | 0.638 | 0.995 | 0.777 | 140:41:16 | — |
| LTRharvest | 131525 | 14358 | 29663 | 539 | 0.484 | 0.964 | 0.644 | 00:21:31 | — |
| LtrDetector | 145789 | 21527 | 29663 | 324 | 0.726 | 0.985 | 0.836 | 06:25:46 | — |
| Total (Including *H. vulgare*) |  |  |  |  |  |  |  |  |  |
| LtrDetector | 322730 | 25882 | 35455 | 496 | 0.730 | 0.981 | 0.837 | 15:23:25 | — |

Aggregate sensitivity is the classification measure in which we saw the most improvement, with our software tool identifying 73.0% of known LTR-RT overall, in comparison to 63.8% by LTRharvest (improvement of 14.4%) and 48.4% by LTRFinder (improvement of 50.8%). Additionally, LtrDetector produced fairly consistent results across the different genomes we tested, ranging from 76.2% on the small genome of *A. thaliana* to 71.2% on the large and repeat-rich*Z. mays*. LtrDetector was the most sensitive tool on the six genomes.

LTR Finder predicted very few false positives and was the most precise tool overall at 99.5%. LtrDetector came in second at 98.1% followed by LTRharvest at 96.4%. All three tools found many more true positives than false positives, resulting in high precision overall.

On the F1 composite measure, LtrDetector again achieves the highest score, outperforming LTR Finder by 7.7% (83.7 vs. 77.7) and LTRharvest by 30% (83.7 vs. 64.4). These results demonstrate that LtrDetector strikes a balance between thorough collection of known LTR-RTs and avoiding spurious detections.

Our evaluation criteria are dependent on the consensus sequences available in Repbase, so we will not be able to definitively classify the majority of proposed LTR-RTs as true positives or false positives. Such detections will be unconfirmed, but could potentially be novel repeats. On the first five genomes (excluding barley), LtrDetector and LTRharvest propose similar amount of retrotransposons —145789 and 131525, respectively. LTR Finder is more conservative with only 88250 guesses.

The total number of detections helps in deriving an estimate of the percentage of a given genome that is composed of full-length LTR-RTs. We summed the length of each detection in base pairs and divided this total by the number of base pairs in the entire genome. This produced estimates of 6.64% LTR-RT content for*A. thaliana*, 14.86% for*O. Sativa*, 15.86 % for*G. max*, 20.22 % for*H.Vulgare*, 28.66 % for*S. bicolor*, and 43.77% for*Z. mays*.

The three tools have vastly different run times. LtrDetector works in parallel by default, whereas we had to configure the other two tools to process several chromosomes simultaneously using the GNU parallel command utility.

The experiments were run on all four cores of an Intel i5 machine with 16 GB RAM running Ubuntu. We recorded wall-clock time using the Linux time command. LTRharvest was by far the fastest tool, capable of processing the five smallest genomes in 21 minutes and 31 seconds. On the other end of the spectrum, LTR Finder took almost 141 hours — about 6 days. LtrDetector’s runtime efficiency was in the middle (around 6 hours and 30 min). LTRharvest and LTR Finder were unable to process the*Hordeum vulgare* genome, whereas LtrDetector was able to do so in about 9 hours.

LtrDetector and LTR Finder generally had the highest memory requirements of the the three tools. LtrDetector nearly maxes out the 16 GB of RAM on the barley genome. LTR Finder requires much more memory than the other tools on the maize genome. LTRharvest uses less memory overall, although memory constraints may be contributing to its failure on the barley genome.

The above experiments suggest that LtrDetector substantially advances the state of the art for finding LTR-RT elements*de-novo*. In comparison to older software tools, it delivers more accurate predictions in reasonable time using memory readily available on modern personal computers. Its capabilities are proven not only on simple model organisms but also on a wide variety of plant genomes.

Crucially for researchers, the tool is easy to install and run and will perform well on an ordinary desktop computer. It provides a robust set of default parameters for maximum generality, but still allows for user configuration via command-line options. As more genome sequences become available, the utility of tools like LtrDector will only increase.

### Post-processing manual annotation aid

LtrDetector is unique in providing a simple visualization tool to aid with manual verification of putative LTR-RTs.

For each LTR-RT identified by LtrDetecor, the script will produce a colorful graph showing distances between k-mers (short words of length k) and their nearest copies as well as markers for the start and end locations of each LTR. This signal will ideally show two flat plateaus representing two LTRs. More details are given in the Methods Section.

### Comparisons to related tools

LtrDetector represents an innovative approach to repeat detection that differs greatly from its predecessor tools.

LtrDetector depends on signal processing techniques. The use of a signal of k-mer distances as an indication of repeat locations is the first of its kind. Both of our closest competitor tools use completely different approaches utilizing suffix-arrays, which are complex data structures that have been widely used in text processing [54].

LTRharvest uses a suffix-array to detect initial maximal repeats — seeds — and a greedy dynamic programming algorithm called X-drop extension to expand from the seeds [50]. It can filter based on length, LTR identity, target site duplications and the palindromic LTR motif (i.e. TG..CA box).

LTR Finder begins with all sets of exact repeats found by the suffix-array [52]. Each member in every set is considered in a pair-wise fashion. The region between the two start coordinates in a pair is aligned with the region between the two end coordinates. The pairs are merged if the alignment is above a certain threshold.

Similar to LtrDetector, LTR Finder uses the Smith-Waterman local alignment algorithm [55] for boundary adjustment. LTR Finder concludes with an aggressive filtering system based on searching for target site duplications, the TG..CA box, primary binding sites, and the proper protein domains in the interior sequence.

### Future work

Future work will seek to improve time efficiency, largely by reducing our dependence on local alignment algorithms, which are very slow on longer sequences. The current model uses a high value (2000 bp) for the maximum LTR length parameter, which produces excessively long alignment windows in most cases.

One approach could be to dynamically set the size of the alignment window based on the initial LTR length. Another approach would be to substitute the Smith-Waterman algorithm with more efficient approximation algorithms. We will also attempt to reduce memory consumption by optimizing the C++ code-base.

## 3 Methods

### Overview

We used a variety of computational techniques to perform*de-novo* signature-based detection of Long terminal repeat (LTR) retrotransposons (LTR-RTs). Signature-based tools rely strictly on specific structural features of LTR-RTs, e.g. the presence of two flanking LTRs, without referring to the nucleotide sequences of known elements. The main contribution of this study is a software package called LtrDetector. The tool utilizes methods from signal processing, using the distances between copies of k-mers — short nucleotide sequences of length k — to determine the location of LTRs.

At a high level, LtrDetector locates LTR-RTs using the following steps (Figure 1):

- Mapping each nucleotide in a sequence to a positive or negative numerical score recording the distance to the closest exact copy of the k-mer starting at that nucleotide.
- Processing the scores to merge adjacent stretches of similar scores (i.e. plateaus).
- Collecting plateaus and pairing those whose distance scores point to each other.
- Correcting LTR coordinates via local alignment of the regions surrounding each plateau in a pair.
- Removing faulty candidates based on sequence identity, element length, and structural similarity to other types of transposable elements (TEs).

**Figure 1.**
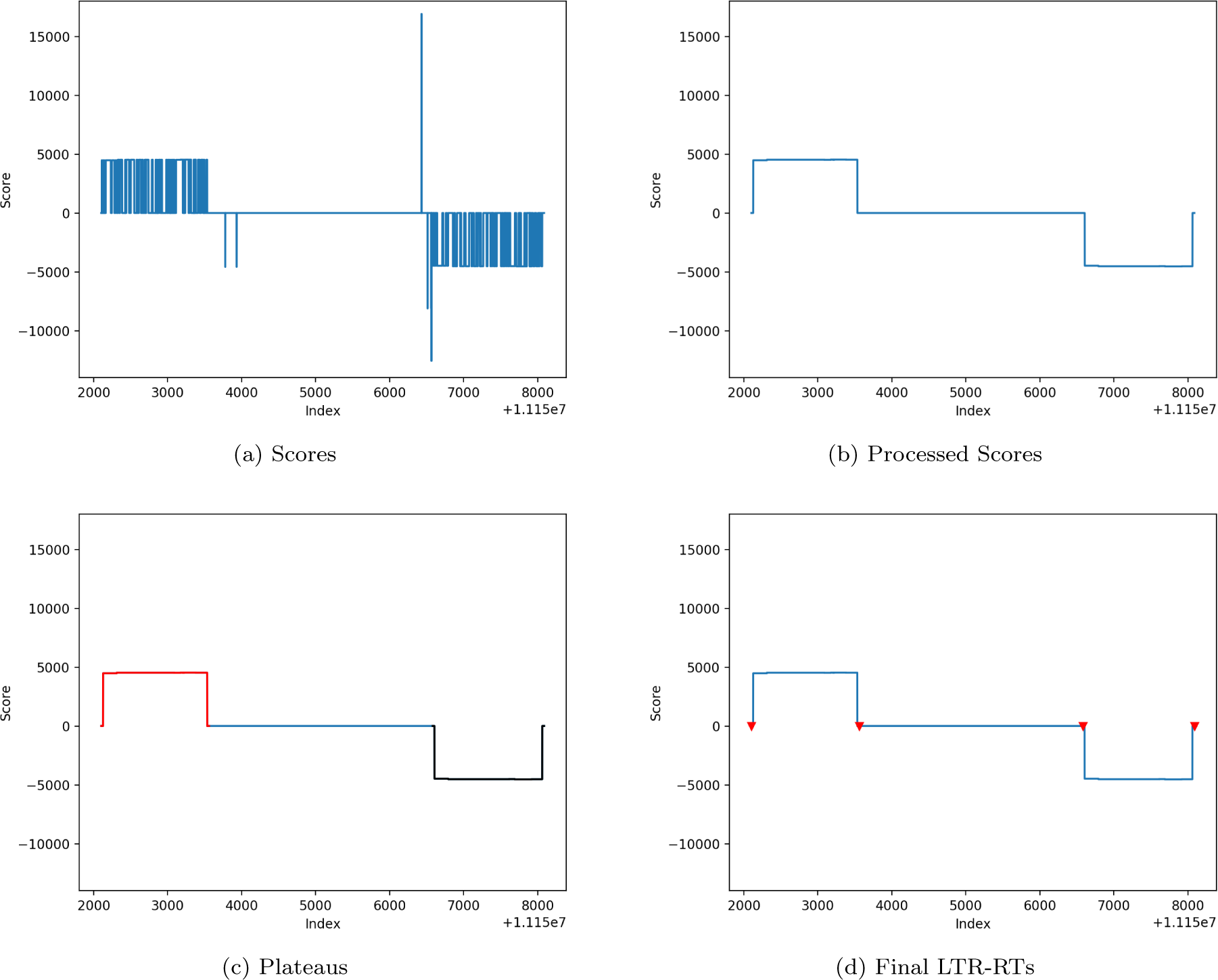
Method overview. LtrDetector is a software tool for locating long terminal repeat (LTR) retrotransposons (RTs). (a) A sequence of scores reflects the distance to the closest exact copy of the k-mer starting at each nucleotide. (b) Smoothed scores are produced after adjacent spikes are merged into a contiguous region. (c) Plateau regions are identified. Separate plateaus here are represented by black and red lines. (d) Plateaus are paired and their boundaries are adjusted. The red triangles denote the start and end coordinates for each LTR.

Scoring the input sequence At the time of insertion into the genome, the two LTRs of an LTR-RT will be identical [4]. Centuries worth of mutation will lead to some degeneration, but the LTRs should retain a high degree of homology.

The goal of the scoring step is to mark the genomic distance to the nearest exact copy of every k-mer in the input sequence using a data-structure called a hash-table. A hash-table is conceptually similar to a dictionary, containing entries mapping a unique key to an associated value. It employs a mathematical function called hashing to convert a key into an index, which is used to look up the value in the hash-table’s underlying array.

LtrDetector utilizes a hashing function specific to DNA sequences. Each nucleotide (A, C, G, and T) in a k-mer is encoded as a digit (0, 1, 2, and 3). This digit sequence is considered as a quaternary (base-4) number and converted to a decimal number (base-10) that indicates an index within the array. Horner’s rule is used for efficiently converting the number from its quaternary to its decimal representation [56]. We have used similar data structures successfully in other software tools [57–60]. For example, the 5-mer ACCTG is transformed to 01132 (base 4) and then to 94 (base 10), mapping it to the 94^*^th^*^ cell in the array. Note that the array for storing all k-mers will be of length 4^*^k^*^.

LtrDetector traverses the input sequence nucleotide by nucleotide, computing the index for the k-mer starting at each position. As it encounters a particular kmer for the first time, it will fill in the hash-table value with the initial location. Whenever that k-mer is found again, we update the hash table with the new location and report the score at that index as the distance between the current copy and the previous copy. Distances are scored both forward and backward in the genome. Scores will be updated if a copy is found closer downstream. The distance between the k-mer and its copy must be within a specific range due to the length properties of LTR retrotransposons [1].

Processing scores The raw scores yielded by the previous step are processed to accentuate meaningful patterns. Wherever there is a significant repeat in the genome, there should be an extended, semi-continuous sequence of similar scores. However, any mutation will cause gaps in these stretches. LtrDetector first identifies all continuous stretches of non-zero scores, categorizing them as “keep” — K — if they are longer than or equal to a minimum seed value, (default: 10 bp), or “delete” — D — if they are not. The forward merging step merges a D section with a neighboring K section if the two are separated by a gap of less than a certain size (default: 200 bp). To merge, the scores belonging to the D section are overwritten with the median score of the adjacent K section, as are the scores in between. This D section is re-categorized as a K. Neighboring K sections will be merged by re-scoring only the gap section, using the median score of one of the two K sections. Next, the backward merging step proceeds in the opposite direction, merging all D sections that appear upstream of K sections and are missed by the forward merging step. After both passes, all remaining D selections are overwritten to zero to reduce noise. We illustrate this procedure in Figure 2.

**Figure 2.**
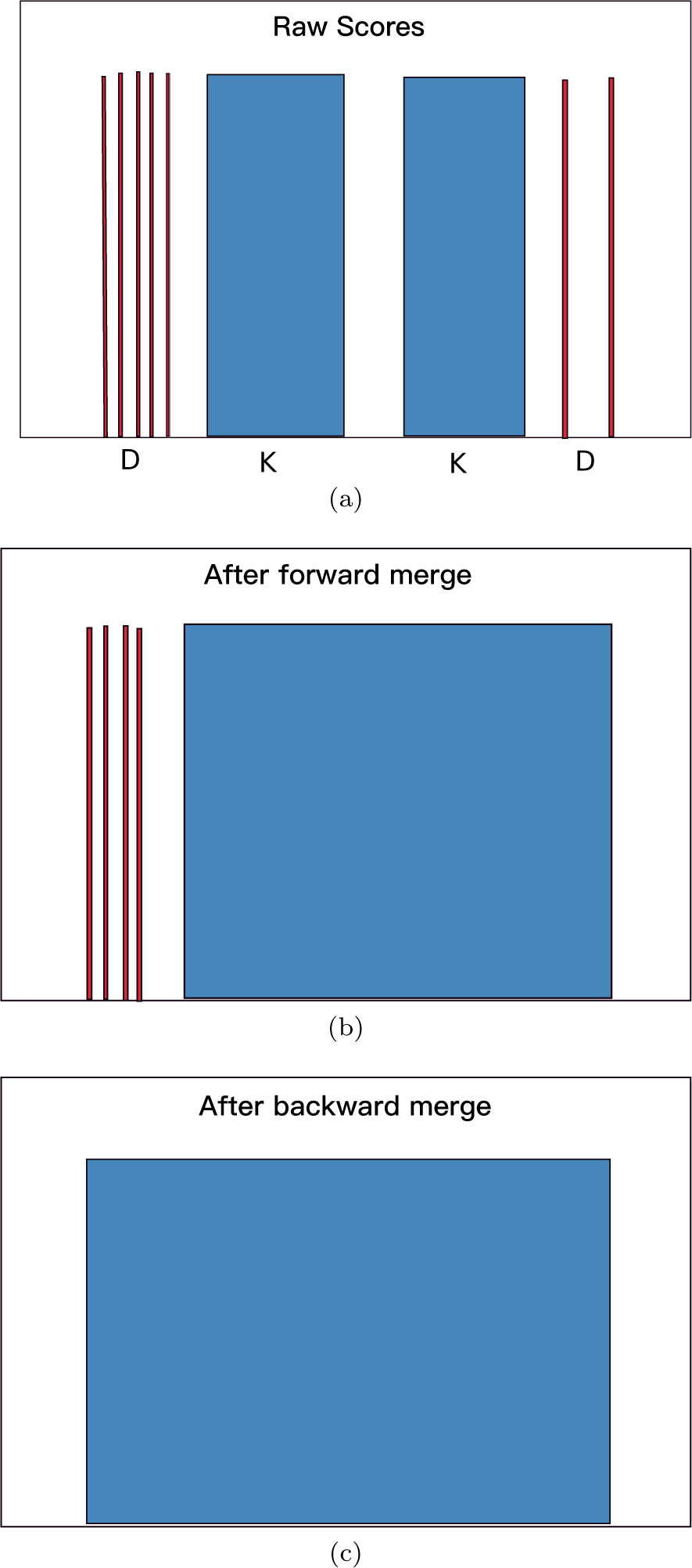
Contiguous stretches of the same non-zero score are identified and marked as keep (K) or delete (D). The forward pass merges K sections toward each other and adjacent D sections. The backward pass merges remaining D sections that are close to K sections.

### Pairing plateaus

The merging step should produce wide plateaus in the scoring signal. The magnitude of their scores can be thought of both as the height of the plateau and as the distance towards its match. The sign of the scores indicate the direction — positive for downstream and negative for upstream. For instance, a plateau of width 200 and height +8000 should imply a similarly wide plateau of height −8000 starting about 8000 base pairs downstream. In this way, the scores of the two plateaus point towards each other.

Another hash-table-like data structure helps pair matching plateaus to form a full retrotransposon. Plateaus are assigned to a bin based on the magnitude of their height, with each bin holding plateaus within a certain range of height. The algorithm then steps through the candidates, placing each positive plateau into the appropriate bin as it is encountered. Each negative plateau will be assigned an initial bin, which will be searched for a positively-scored plateau located at the proper distance; recall that this distance is implied by the height of the negative plateau. Because we allow for some difference in height, the bins immediately above and below the initial one may be inspected. If a match is found, the two regions are returned and listed as a candidate LTR pair. If not, the negative plateau is discarded.

### Boundary correction

The newly paired LTR candidates merely approximate the boundaries of a putative retrotransposon because the sensitivity of k-mers to mutations. Next, we use the Smith-Waterman local alignment algorithm [55] to sharpen the LTR boundaries. Both LTR regions are expanded according to Equations 4 and 5, and then aligned.

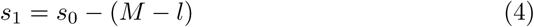

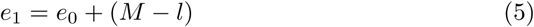

Here,*l*, *s*_0_, and*e*_0_ are the length, the start, and the end of the original region, whereas*s* _1_ and*e* _1_ are the start and end of the expanded region. *M* is the maximum length of an LTR, set at 2000 bp by default. The returned alignment indicates the corrected boundaries for the putative LTR-RT. Alignment identity scores are stored for later use.

### Filtering

Several filters are applied to reduce the number of false positives. LTR identity scores from the previous step are used for discarding all entries whose paired LTRs exhibit sequence similarity below a given threshold (default: 85%). Then elements are filtered by size to remove those where either the LTR or the whole element is too small or too large, using values typical of known LTR-RTs. Our default range for the full element is 2000–18000 and 100–2000 for its LTRs.

Finally, the candidates are analyzed to determine whether they exhibit features of DNA transposons, which are another type of TE that appear in high copy number in many genomes. DNA transposons of the same family can appear in close proximity and be falsely identified as LTR-RTs by the previous steps. DNA transposons contain terminal inverted repeats, meaning that the reverse complement of the beginning sequence appears at the end of the element.

LtrDetector locally aligns the first 100 nucleotides of each LTR with the reverse complement of its last 100 nucleotides. If this resulting alignment is sufficiently long (> 15 bp), the element may represent two DNA transposons within close distance to each other; this element is discarded.

### Reporting

Elements that pass all of the filters are reported as LTR-RTs in BED file format. Each entry contains start and end coordinates for the entire retrotransposon as well as for its LTRs. LtrDetector runs in parallel by default; it can analyze one chromosome on each machine core.

### Ground truth generation

In order to assess the accuracy of our signature-based predictions, we built a pipeline for assembling ground truth using previously known sequences of LTR-RTs. Repbase is the most comprehensive database for repetitive elements, containing consensus sequences for a wide variety of genomes [61]. The Repbase browser system provides FASTA files containing the LTRs and interior sequences for full LTR-RTs separately. The ground truth were constructed using two related, complementary approaches.

In the first approach, we downloaded these files and parsed them to append the LTR sequences before and after their associated interior sequence to form full LTR-RT elements. We then performed a BLAST search for these complete elements against the input genome, processing the output from BLAST to accept only those results that represent 100% coverage of the query as well as 70% or more identity.

In the second approach, we built a pipeline around RepeatMasker — the standard database-driven tool for repeat detection. RepeatMasker also uses Repbase as its default source of repeat consensus sequences.

Instead of concatenating the LTRs and their interior regions before the search, we searched for them separately in RepeatMasker’s output. These entries are used for extracting LTR-RT coordinates by finding two 100%-coverage detections of the same LTR that are 200–12000 bp apart as defined by their start coordinates. The corresponding interior element was required to appear somewhere in between.

The outputs of the two pipelines are merged and duplicates are removed. The reason we use both pipelines is that the results from the RepeatMasker pipeline are dependent upon estimated length parameters, but do a better job finding LTR-RTs with more degenerate interior regions, whereas the BLAST data is free from guesswork but stricter about enforcing the canonical structure of LTR-RTs.

### False positive evaluation

We built a false positive detection pipeline by parsing RepeatMasker output to determine when putative LTR-RTs overlap with non-LTR repeats. Repeat-Masker detections that do not belong to an LTR (excluding simple and low-complexity repeats) are compared with the predicted LTR-RTs. If two repeat detections of the same type overlap by more than 80% with the two supposed LTRs of a putative LTR-RT, this element is considered as a false positive. This approach was inspired by another study for detecting Miniature Inverted-repeat Transposable Elements (MITEs) [62]; in that study, a putative element is considered a false positive if it overlaps with any non-MITE elements. Repbase is by no means a complete record of repetitive elements, so neither the ground truth nor the false positive annotations will be comprehensive. Accordingly, a large amount of detections by LtrDetector will overlap with neither set and will be impossible to evaluate against existing databases. Nonetheless, this approach is, in our opinion, the best available method for evaluating the false positives of tools for detecting LTR-RTs.

### Data

We validated the results of LtrDetector using a variety of genomes. An initial test replicated the experiment in a study by Lerat [1], testing multiple tools on the X Chromosome of the*D. melanogaster* (Dm3) against a ground-truth annotation assembled from RepeatMasker. We performed similar analysis on the following genomes:

- *Arabidopsis thaliana* (TAIR10):http://plants.ensembl.org/Arabidopsisthaliana/Info/Index
- *Hordeum vulgare* (HvIbscPgsbV2):http://plants.ensembl.org/Hordeumvulgare/Info/Index
- *Oryza sativa Japonica* (IRGSP1):http://plants.ensembl.org/Oryzasativa/Info/Index
- *Sorghum bicolor* (SorghumBicolorV2):http://plants.ensembl.org/Sorghumbicolor/Info/Index
- *Zea mays* (ZeaMaysAGPv4):http://ensembl.gramene.org/Zeamays/Info/Index
- *Glycine max* (Gmax 109):http://www.plantgdb.org/XGDB/phplib/download.php?GDB=Gm

## Availability and requirements

The source code (C++ and Python) is available as Additional file 1.

**Project name:** LtrDetector.

**Project home page:** http://github.com/TulsaBioinformaticsToolsmith/LtrDetector:

**Operating system(s):** UNIX/Linux/Mac.

**Programming language:** C++ and Python.

**Other requirements:** BLAST https://blast.ncbi.nlm.nih.gov/Blast.cgi(https://blast.ncbi.nlm.nih.gov/Blast.cgi) and Bedtools http://bedtools.readthedocs.io/en/latest/(http://bedtools.readthedocs.io/en/latest/).

**Python:** NumPy, Matplotlib, Pandas.

**License:** Creative Commons license (attribution + non-commercial + no derivative works). Any restrictions to use by non-academics: License needed.

## Declarations

### Ethics approval and consent to participate

Not applicable.

### Consent to publish

Not applicable.

### Availability of data and materials

The source code of LtrDetector, analysis scripts, and long terminal repeats retrotransposons are available as Additional files 1–9.

### Author’s contributions

HZG designed the software and the case studies, implemented the scoring module, and wrote the manuscript. JDV implemented the software and the evaluation pipelines, conducted the experiments, produced the results, and wrote the manuscript.

**List of Abbreviations**
Bp: base pairs
TE: transposable element
LTR: long terminal repeat
RT: retrotransposon
LTR-RT: long terminal repeat retrotransposon.

## Acknowledgements

Not applicable.

## Additional Files

Additional file 1 — The LtrDetector Software This compressed file (.tar.gz) includes the C++ source code of LtrDetector.

Additional file 2 — Scripts for evaluation pipeline and visualization This compressed file (.tar.gz) includes the Python code for the evaluation pipeline and visualization scripts.

Additional file 3 — The synthetic sequences This compressed file (.tar.gz) includes the synthetic sequences with different mutation rates.

Additional file 4 — Long Terminal Repeat (LTR) retrotransposons found by LtrDetector in *Arabidopsis thaliana* This compressed file (.tar.gz) includes the LTR retrotransposons found by LtrDetector in BED format.

Additional file 5 — LTR retrotransposons found by LtrDetector in*Glycine max* This compressed file (.tar.gz) includes the LTR retrotransposons found by LtrDetector in BED format.

Additional file 6 — LTR retrotransposons found by LtrDetector in*Hordeum vulgare* This compressed file (.tar.gz) includes the LTR retrotransposons found by LtrDetector in BED format.

Additional file 7 — LTR retrotransposons found by LtrDetector in*Oryza sativa Japonica* This compressed file (.tar.gz) includes the LTR retrotransposons found by LtrDetector in BED format.

Additional file 8 — LTR retrotransposons found by LtrDetector in*Sorghum bicolor* This compressed file (.tar.gz) includes the LTR retrotransposons found by LtrDetector in BED format.

Additional file 9 — LTR retrotransposons found by LtrDetector in *Zea mays* This compressed file (.tar.gz) includes the LTR retrotransposons found by LtrDetector in BED format.

